# Isolation and Bioassay of Linear Veraguamides from a Marine Cyanobacterium (*Okeania sp.*)

**DOI:** 10.1101/2025.01.18.633713

**Authors:** Stacy-Ann J. Parker, Andrea Hough, Thomas Wright, Neil Lax, Asef Faruk, Christian K. Fofie, Rebekah Simcik, Jane E. Cavanaugh, Benedict J. Kolber, Kevin J. Tidgewell

## Abstract

Marine cyanobacteria have gained momentum in recent years as a source of novel bioactive small molecules. This paper describes the structure elucidation and pharmacological evaluation of two new, veraguamide O (**1**) and veraguamide P (**2**), and one known, veraguamide C(**3**), analogs isolated from a cyanobacterial collection made in the Las Perlas Archipelago of Panama. We hypothesized that these compounds would be cytotoxic in cancer cell lines. The compounds were screened against HEK-293, estrogen receptor positive (MCF-7), and triple-negative breast cancer (MDA-MB-231) cells as well as against a broad panel of membrane bound receptors. The planar structures were determined based on NMR and MS data along with comparison to previously isolated veraguamide analogs. Phylogenetic analysis of the collection suggests it to be an *Okeania* sp., a similar species to the cyanobacterium reported to produce other veraguamides. Veraguamide O shows no cytotoxicity (greater than 100 M) against ER positive cells (MCF-7) with 13 M IC_50_ against MDA-MB-231 TNBC cells. Interestingly, these compounds show affinity for the sigma2/TMEM-97 receptor making them potential leads for development of non-toxic sigma 2 targeting ligands.

## Introduction

Natural products are crucial sources for drug development and discovery. In the last two decades, marine cyanobacteria have emerged as a leading source for novel bioactive natural products. The high degree of chemical diversity among cyanobacterial natural products is attributed to their biosynthetic machinery, which integrates polyketide synthases (PKS) and nonribosomal peptide synthases (NRPS) into mixed biosynthetic pathways resulting in diverse bioactive secondary metabolites. This broad spectrum of marine cyanobacterial natural products often translates into a wide array of biological activities, including but not limited to, anticancer,^1^ antibacterial,^2^ antimalarial,^3^ antiinflammatory,^4^ and neuroactive^5-7^ activities.

The potential for cyanobacteria as a source for novel antitumor cytotoxicity is particularly exciting. The goal of this study was to isolate and identify new molecules from marine cyanobacteria with potential antitumor activity. We report here on the isolation and characterization of one known and two novel natural products within the veraguamide family. Veraguamides have previously been shown to have cytotoxic effects in the human lung cancer cell line H-460.^8^ Both cyclic and linear veraguamides have been reported with cyclic versions being more common. Interestingly, the majority of characterized marine cyanobacteria cytotoxic compounds are cyclic depsipeptides, and some of the most potent activities were observed against H-460 lung cancer cell lines. ^9, 10-12^

Herein, we report on the isolation and characterization of new linear hexadepsipeptides, veraguamides O (**1)** and P (**2)**, along with the known cyclic compound veraguamide C (**3**) ^8, 13^ from a Panamanian marine cyanobacterium. While depsipeptides have been screened against human lung (H-460), colon (HCT-116), and breast cancer (MCF-7) cell lines, so far, none have been investigated for their cytotoxic effects against the more resistant and aggressive triple-negative breast cancer (TNBC) cells. Furthermore, the cytotoxic effects for most linear depsipeptides against human cancer cell lines have not been determined. ^14 8^

TNBC represents 15-20% of breast cancers and is characterized by a lack of estrogen receptors, progesterone receptors, and human epidermal growth factor receptor 2 (HER2) receptors.^15^ Treatment options for TNBCs are primarily limited to chemotherapy and radiation or surgical resection of early stage disease.^16^ This is in contrast to hormone receptor positive breast cancers, which are successfully targeted by small molecules or antibodies that specifically target these receptors (e.g. estrogen receptors)^17^. Due to the limited treatment options, there is a need to develop new leads for TNBC treatment.

We tested the hypothesis that our isolated veraguamides would produce cytotoxic effects including in TNBCs. We tested this hypothesis in HEK-293 (non-cancerous), MCF-7 (ER positive), and MDA-MB-231 (TNBC) cell lines. We also evaluated the potential of these molecules to be overtly toxic in wildtype mice.

## Results and Discussion

A cyanobacterial mat (**Figure 3A**) was collected off the coast of Isla Mina in the La Perlas Archipelago, Panama (GPS coordinates: N 8 29.717 W 78 59.947) and given the extraction ID of DUQ0008. This collection was extracted with CH_2_Cl_2_-MeOH (2:1) to afford 3.3 g of crude extract. This crude extract was fractionated by silica gel chromatography to yield nine fractions labeled A (100% hexanes) through I (100% methanol). Fraction I, eluted with 100% methanol, was further purified using accelerated chromatography and reversed-phase HPLC to afford **1** (3.8 mg) and **2** (0.8 mg) (**Figure 1**); Fraction G, eluted with 10% hexanes/90% methanol, was further purified using accelerated chromatography and reverse-phase HPLC to afford **3** (1.7 mg) (**Figure 1**). All were recovered as colorless oils.

**Figure 1:**
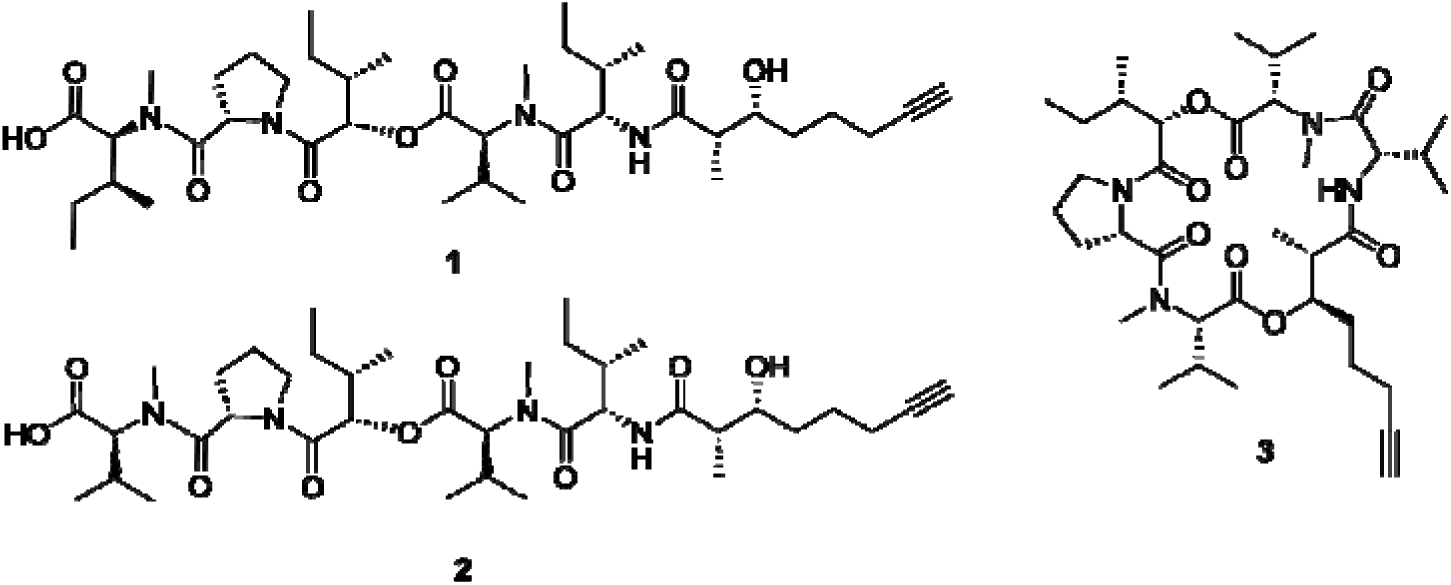
Structures of isolated veraguamides O (1), P (2), and C (3).

HRESIMS of **1** in positive mode provided molecular ion peaks at m/z 735.4804 [M + H], and 757.4626 [M + Na] suggesting a molecular formula of C_39_H_66_N_4_O_9_, corresponding to nine degrees of unsaturation. The structure of **1** was established by analysis and correlations of peaks in the NMR spectra which revealed that it was a depsipeptide (**Table 1**), with α-protons (δ_H_ 4.67–4.87), an exchangeable amide (NH) proton (δ_H_ 6.82), two *N*-methyl groups (δ_H_ 3.11 and 3.06), and deshielded signals in both the ^1^H and ^13^C NMR spectra typical of oxymethines (δ_H/C_ 4.95/76.5 and 3.60/71.7) one of which was adjacent to an ester linkage. This observation was further supported by ^13^C NMR resonances derived from HSQC and HMBC experiments, assignable to six ester, amide, and carboxylic acid carbonyls (δ_C_ 175.6, 173.1, 170.3 and 170.1). The aforementioned together with the terminal alkyne (δ_C_ 84.4, 68.7, δ_H_ 2.16) of the 3-hydroxy-2-methyloctynoic acid (Hmoya) residue accounted for eight of the nine degrees of unsaturation and suggested that **1** possesses an additional ring. Further analysis of the COSY, TOCSY, HSQC and HMBC data for **1** in CD_3_CN (**Table 1**, **Figure 2**, and Supporting Information **Figures S1-S6**) revealed the presence of four proteinogenic amino acid residues (*N*-Me-Ile, Pro, *N*-Me-Val and Ile) and the hydroxy acid, 2-hydroxy-3-methylpentanoic acid (hmpa). Given that the proline residue accounted for the final degree of unsaturation, and that there are two terminal groups (*N*-Me-Ile and Hmoya), it was clear that the compound is linear. The fragments *N*-Me-Ile–Pro–Hmpa–*N*-Me-Val–Ile–Hmoya were readily constructed with the help of TOCSY and HMBC data.

**Figure 2.**
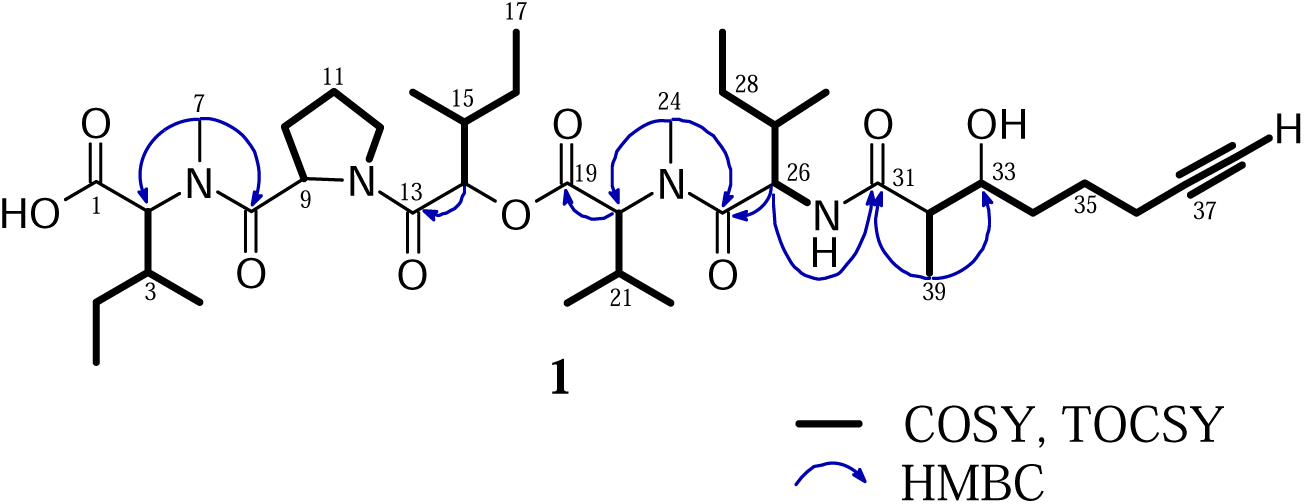
COSY and TOCSY connectivity and selected HMBC correlations for **1**.

**Table 1.** NMR spectroscopic data for veraguamides O (**1**) and P (**2**) in CD_3_CN.

| **1** | **2** |
| --- | --- |

**Table 1.** NMR spectroscopic data for veraguamides O (**1**) and P (**2**) in CD<sub>3</sub>CN
| Residue | Pos. | $\delta_C$, type | $\delta_H$ (J in Hz) | HMBC<sup>a</sup> | $\delta_C$ | $\delta_H$ (J in Hz) | HMBC<sup>a</sup> |
| --- | --- | --- | --- | --- | --- | --- | --- |
| *N*-Me-Ile<sup>d</sup>/<br>*N*-Me-Val<sup>e</sup>-<br>1 | 1. | 170.3, C |  |  | 170.3 |  |  |
|  | 2. | 61.5, CH | 4.67, d (10.7) | 7 | 61.5 | 4.70 |  |
|  | 3. | 32.6, CH | 2.01, m |  | 27.1 | 2.23 |  |
|  | 4. | 24.6, CH<sub>2</sub> | 1.15 |  | - | - |  |
|  | 5. | 15.1, CH<sub>3</sub> | 0.97 | 2, 3, 4 | 19.0 | 0.90 | 2, 3, 6 |
|  | 6. | 9.9, CH<sub>3</sub> | 0.87 | 3, 4 | 18.1 | 0.80 | 2, 3, 5 |
|  | 7. | 31.7, CH<sub>3</sub>, | 3.06, s | 2, 8 | 31.6 | 3.06, s | 8 |
| Pro | 8. | 173.1, C |  |  | 172.9 |  |  |
|  | 9. | 56.8, CH | 4.79, q (8.1, 5.2) |  | 56.7 | 4.80 |  |
|  | 10. | 28.2, CH<sub>2</sub> | 1.76, m |  | 28.2 | 1.79, m |  |
|  | 11. | 24.5, CH<sub>2</sub> | 2.23, 2.09 |  | 24.5 | 2.22, 2.08 |  |
|  | 12. | 47.2, CH<sub>2</sub> | 3.83, m<br>3.60<sup>c</sup> |  | 47.0 | 3.82, m<br>3.61<sup>c</sup> |  |
| Hmpa | 13. | 170.3, C |  |  | 170.3 |  |  |
|  | 14. | 76.5, CH | 4.95 | 13, 15, 16,<br>18 | 76.2 | 4.90 |  |
|  | 15. | 36.2, CH | 1.93<sup>b</sup> |  | 36.0 | 1.93<sup>b</sup> |  |
|  | 16. | 24.1, CH<sub>2</sub> | 1.67, s |  | 23.8 | 1.67 |  |
|  | 17. | 10.4, CH<sub>3</sub> | 0.89 | 15, 16 | 10.3 | 0.89 | 15, 16 |
|  | 18. | 14.0, CH<sub>3</sub> | 1.00 | 14, 15, 16 | 13.8 | 0.99 | 14, 15, 16 |
| *N*-Me-Val-2 | 19. | 170.1, C |  |  | 170.3 |  |  |
|  | 20. | 61.7, CH | 4.87, d (10.7) | 19, 21, 23,<br>24 | 61.6 | 4.86, d (10.6) | 19, 21 |
|  | 21. | 27.1, CH | 2.23 |  | 27.1 | 2.26 |  |
|  | 22. | 19.1, CH<sub>3</sub> | 0.99 | 20, 21, 23 | 19.0 | 0.99 | 20, 21, 23 |
|  | 23. | 17.9, CH<sub>3</sub> | 0.79, d (6.7) | 20, 21, 22 | 18.1 | 0.79 | 20, 21, 23 |
|  | 24. | 31.3, CH<sub>3</sub> | 3.11, s | 20, 25 | 31.4 | 3.12 | 20 |
| Ile | 25. | 173.1, C |  |  | 173.0 |  |  |
|  | 26. | 52.9, CH | 4.70<sup>c</sup> | 25, 27, 31 | 52.8 | 4.70, m |  |
|  | 27. | 36.9, CH | 1.81, m |  | 36.8 | 1.83, m |  |
|  | 28. | 24.5, CH<sub>2</sub> | 1.57, 1.15 |  | 24.5 | 1.59, 1.17 |  |
|  | 29. | 10.4, CH<sub>3</sub> | 0.90 |  | 10.3 | 0.90 | 27, 28 |
|  | 30. | 14.9, CH<sub>3</sub> | 0.90 | 26 | 14.9 | 0.91 | 26, 27, 28 |
| Hmoya | 31. | 175.6, C |  |  | 175.5 |  |  |
|  | 32. | 45.4, CH | 2.35, m | 31, 33 | 45.1 | 2.34, m | 31, 39 |
|  | 33. | 71.7, CH | 3.60, m |  | 71.6 | 3.61, m |  |
|  | 34. | 32.8, CH<sub>2</sub> | 1.42 |  | 32.7 | 1.41 |  |
|  | 35. | 24.6, CH<sub>2</sub> | 1.67, 1.48 |  | 24.9 | 1.67, 1.47 |  |
|  | 36. | 17.9, CH<sub>2</sub> | 2.19 | 34, 35, 37 | 17.7 | 2.18 | 34, 35, 37,<br>38 |
|  | 37. | 84.4, C |  |  | 84.5 |  |  |
|  | 38. | 68.7, CH | 2.16<sup>b</sup> | 37 | 68.7 | 2.16<sup>b</sup> |  |
|  | 39. | 12.1, CH<sub>3</sub> | 1.09, d (6.6) | 31, 32, 33 | 11.9 | 1.10, d (6.6) | 31, 32, 33 |
|  |  | NH | 6.82, d (9.0) |  |  | 6.81, d (8.6) |  |
|  |  | OH | 3.43 | 34 |  |  |  |

<sup>a</sup>Carbon correlating to proton shift, <sup>b</sup>Signal obscured, <sup>c</sup>Signal partially obscured, <sup>d</sup>Pertaining to **1**, <sup>e</sup>Pertaining to **2**

The continuous sequence of the residues was determined by long-range HMBC connectivities (**Figure 2**) and mass spectral analysis of the partial hydrolysis product of **1**. ESI-MS analysis of the partial hydrolysis product of **1** (**Scheme 1**) provided evidence for molecular ions at m/z 379.3 [M + Na], 355.3 [M - H] and, 419.3 [M + Na] and 395.3 [M - H]. These corresponded to fragments with molecular formulas C_18_H_32_N_2_O_5_ (**5**) and C_21_H_36_N_2_O_5_ (**7**), respectively. The connectivity between the *N*-Me-Ile and Pro residues was established by HMBC correlations between the hydrogens of the *N-*Me (δ 3.06) and the carbonyl carbon at C-8 (δ 173.1), indicating an amide linkage. These correlations together with information derived from the mass of fragment **5**, suggest that the hmpa residue was linked to the proline containing dipeptide fragment. The connection between hmpa and *N*-Me-Val was inferred from ester hydrolysis of **1**. Next, HMBC cross-peaks were observed between the *N-*Me hydrogens of *N*-Me-Val (δ 3.11) and C-25 (δ 173.1) of Ile. Lastly, the α-hydrogen at H-26 (δ 4.70) linked the latter residue to the terminal Hmoya unit through an HMBC connectivity to the carbonyl carbon at δ 175.6 (C-31).

**Scheme 1.**
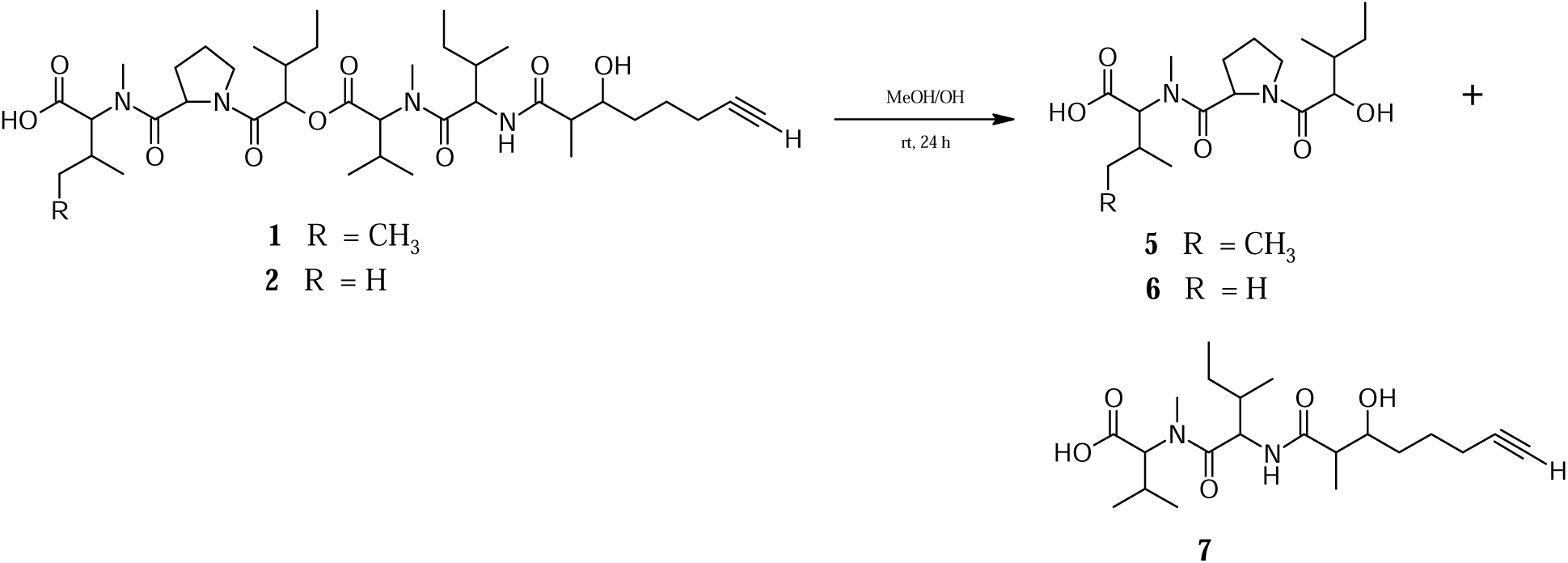
Partial hydrolysis of veraguamides O (**1**) and P (**2**).

Compound **2** (**Figure 1**) also a depsipeptide as suggested by its ^1^H NMR spectrum, displayed molecular ion peaks that indicated the molecular formula C_38_H_64_N_4_O_9_ that also corresponded to nine degrees of unsaturation (Supporting Information **Figures S7-S12**). The six residues, which accounted for all nine degrees of unsaturation, were readily identifiable from characteristic ^1^H and ^13^C NMR resonances and from TOCSY and HMBC correlations (**Table 1**). These data together with the molecular formula, provided evidence for a 3-hydroxy-2-methyloctynoic acid (Hmoya), a 2-hydroxy-3-methylpentanoic acid (hmpa) and four α-amino acid residues (two *N*-MeVal, Pro and Ile). Combining the molecular formula and NMR data for **2** suggested that it was related to **1**, differing by 14 mass units, and provides the justification for its linear structure. The difference of 14 mass units could be explained by the loss of a CH_2_ unit. The loss of the CH_2_ unit likely occurred at the pendant alkyl groups of proteogenic or hydroxy acid side chain. This is supported by (1) the presence of chemical shifts for two *N-*Me (δ_H_/δ_C_ 3.11/31.3 and 3.06/31.7), COSY correlations of the sequence H–39 — H–32 — H–33 — H–34 — H–35— H–36— H–38, which accounted for all three methylene groups of the terminal Hmoya residue and (2) the fact that there are no additional chemical shifts for an amide proton (NH). The position of the CH_2_ loss was confirmed by mass analysis of the partial hydrolysis product of **2** (**Scheme 1**) which, like **1**, provided evidence for fragment **7** (m/z 419.3 [M + Na] and 395.3 [M - H]). This suggested that either the hmpa or *N*-Me-Ile was replaced by 2-hydroxy-3-methylbutanoic acid (hmba) or *N*-Me-Val-1 residue, respectively. A TOCSY experiment readily linked the α-proton (δ 4.90) signal to a spin network belonging to a hmpa residue. HMBC cross-peaks from the hydrogens of the gem-dimethyl group to each other and to the α-carbon at C-2 (δ 61.5) established that the *N*-Me-Ile residue is replaced by the *N*-Me-Val-1 in **2** (**Table 1**).

Compound 3 (**Figure 1**) was isolated as a colorless oil after purification by HPLC and accelerated chromatography from the G fraction of the DUQ0008 extract. Comparison with data from the literature revealed that 3 is veraguamide C, originally isolated from *Oscillatoria margaritifera* collected in Panama.^8, 13^ The six residues identified in 1 and 2, and in particular the Hmoya unit, are consistent with linear and cyclo-hexadepsipetides of the veraguamide series of compounds. These observations together with the recovery of veraguamide C (3) from fraction G (a less polar fraction to I) in this study, strongly suggested that 1 and 2 are related to the veraguamides. Comparative analysis of the residue sequence of these linear depsipeptides with the veraguamides and, other members of the kulolide family^11^ indicated that the compounds are new. 1 and 2 were assigned the trivial names veraguamide O and veraguamide P, respectively.

The relative configuration illustrated in 1 (and 2) was assigned from comparison of the ^13^C NMR data to those of veraguamides A, C (3) and the linear depsipeptide veraguamide L^8^. The absolute configuration of the α-amino acid and hydroxy acid residues in veraguamide A was independently established by Mevers et al.^8^ and Salvador et al.,^13^ revealing only _L_-amino acid residues, and _L_-hmpa and 2*S*, 3*R*-Br-Hmoya after Marfey’s reagent derivatization and formation of esters, respectively. The relative configuration of the residues in the remaining veraguamides (B-G) were deduced based on their virtually identical ^13^C NMR shifts to veraguamide A, and their specific rotations. Based on our analysis and the literature, we expect that these veraguamides also contain the same stereo-configuration as previously reported analogs; however we chose to utilize material for pharmacological testing rather than confirmation of stereoconfiguration.

### Morphological and Phylogenetic Analysis of DUQ0008

The DUQ0008 collection was a dark green/grey marine cyanobacterial mat collected in ∼1 m of water off the coast of Isla Mina in the Las Perlas Archipelago, Panama (**Figure 3A**). Confocal (**Figure 3B**) and transmitted light (**Figure 3C**) microscopy was performed in order to visualize the microscopic structure of the organism. Based on the morphology of the specimen, this sample was field identified as belonging to the genera of *Oscillatoria* or *Moorea*. In order to genetically determine the species of marine cyanobacteria, sequencing of the 16S rRNA gene was performed. The DUQ0008 16S rRNA sequence was first compared in an unbiased fashion using the NCBI Standard Nucleotide BLAST.^18^ The top identified hit with this search (99% alignment) was a species in the *Okeania* genus that produces a cytotoxic cyclohexadepsipeptide, odoamide.^19, 20^ We next built a phylogenetic tree using a collection of sequences from Engene et al that highlights the relatedness of natural product producing marine cyanobacterial strains.^21^ In this tree, we found that DUQ0008 groups closest with an *Okeania* species that produces veraguamides A to C and H to L (**Figure 3D**).^8^ This relationship is consistent with our compound analysis above since DUQ0008 also produced veraguamide C. One of the previously collected veraguamide producers^8^ and our collection both came from the Pacific side of Panama, indicating a possible shared common ancestor of these species. Future collections in this geographic location may yield additional *Okeania* species with novel veraguamide analogs.

**Figure 3.**
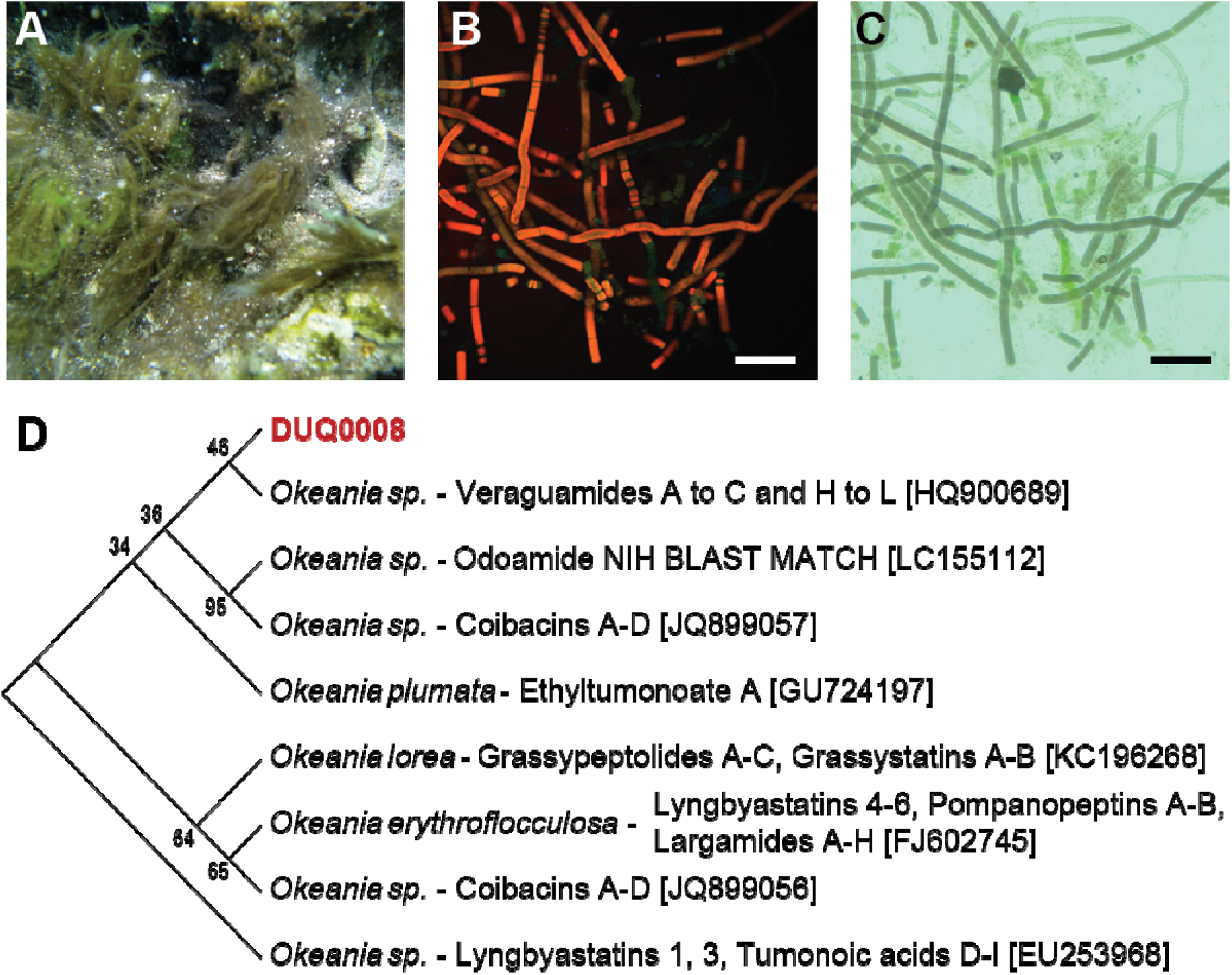
Photos of marine cyanobacteria DUQ0008 and phylogeny. (A) Macroscopic field picture of sample DUQ0008 before collection. (B) Confocal microscopy image of DUQ0008 at 10X magnification with FITC (487 nm), TRITC (560nm) and Cy5 (637 nm) channels. Scale bar = 200 μm. (C) Transmitted light image of DUQ0008. Scale bar = 200 μm. (D) Phylogenetic tree showing the closest related species to DUQ0008 based off 16S rRNA sequencing with comparison to natural product producing cyanobacteria from the *Okeania* genus. Compounds produced by species are indicated if available; accession numbers are in brackets; numbers next to tree show percent of trees showing alignment in 600 replicates.

### Analysis of Veraguamides O, P, and C in Breast Cancer and HEK-293 Cells

Next, we tested the hypotheses that these isolated veraguamides would produce cytotoxic effects in cell lines. Veraguamides O (**1**), P (**2**), and C (**3**), were assessed for their effects against human MCF-7 (estrogen positive), MDA-MB-231 (TNBC) breast cancer, and/or HEK-293 (human embryonic kidney) cell lines. The linear depsipeptide veraguamide O (**1**) displayed selective inhibitory effects against TNBC (MDA-MB-231) cells compared to estrogen positive (MCF-7) breast cancer cells using a MTT assay (**Figure 4A-B**). Veraguamide O (**1**) decreased viability in MDA-MB-231 cells in a concentration dependent manner from 0.14 µM (86% viability) to 14 µM (43% viability) and the IC_50_ of **1** was determined to be 13 ± 4 µM. Functional results from the MTT assay were confirmed microscopically, where veraguamide O (**1**) appeared to induce cell death in TNBC cells (MDA-MB-231) at 10 µM (*data not shown*). The cyclic depsispeptide veraguamide C (**3**) exhibited similar results in both cell lines causing cytotoxicity of 53% in TNBC cells and 65% in MCF7 cells at 15 µM. Values for veraguamide C (**3**) against MCF-7 and MDA-MB-231 cell lines are comparable to those reported in other cell lines previously.^8, 13^ In HEK-293 cells, veraguamide C showed no toxicity up to 10 µM (**Figure 4C**). While these values do not rise to a level to be considered strong cytotoxic leads, it is interesting to note the difference in activity between the cyclic and linear analogs. To our knowledge, maedamide is the only linear depsipeptide that exhibits antitumor cytotoxicity in cell culture.^14^ Given the low potential for developiong these as cytotoxic agents, we decided to look into potential binding to membrane bound receptors for the possibility that the differential activity was due to a modulation of the cells through a specific receptor.

**Figure 4.**
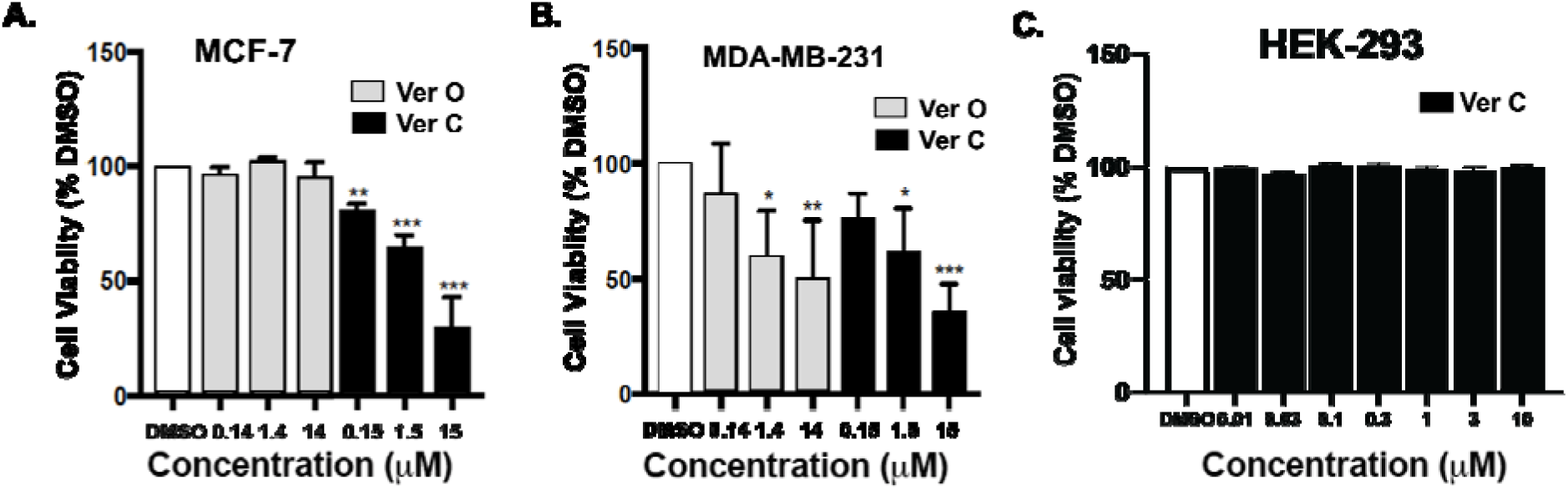
Veraguamides O and C selectively reduce viability in estrogen receptor positive and TNBC cells. Isolated veraguamides O or C activity (A) in the estrogen positive breast cancer cell line, MCF-7 and (B) in the TNBC cell line, MDA-MB-231. Data represent the mean value +/- SE of three different experiments. (C) Veraguamide C has no effect on cell viability in HEK-293 cells. *P<0.05, **P<0.01, ***P<0.001; **** p<0.0001 vs control group determined by one-way ANOVA with the Bonferroni post hoc test.

### Receptor binding affinity

The parent fractions (fractions G and I) as well as pure natural products **1** and **3**, were screened against membrane bound receptors by the Psychoactive Drug Screening Program (PDSP)(**Table 2**). The compounds tested showed only strong binding to the simga2/TMEM97 receptor. Veraguamide C (**3**) was found to bind with an IC_50_ of 320 nM, while veraguamide O (**1**) showed an IC_50_ of 720 nM. This affinity for the sigma 2/TMEM97 receptor is interesting in light of the cancer cell data because this target has been shown to be upregulated and important in activity against MDA-MB-231 cells.^22^ This target has also been shown to be involved in activation of the Store Operated Calcium Entry (SOCE) pathway^23^ within MDA-MB-231 cells leading to apoptosis.

**Table 2.** Binding affinity results for fractions and pure compounds against the Sigma 2 receptor (Transmembrane Protein 97 (TMEM97)).

|  | Primary:<br>Percent Inhibition of Binding | Secondary:<br>IC<sub>50</sub> |
| --- | --- | --- |
| Fraction G | 60% | 1085 ng/mL |
| Fraction H | 54% | 1874 ng/mL |
| Fraction I | 68% | 424 ng/mL |
| Veraguamide C (**3**) | 65% | 316 nM |
| Veraguamide O (**1**) | 87% | 721 nM |

*\* Fractions and compounds that pass primary binding criterion of greater than 50% inhibition at 10 μM are tagged for secondary radioligand binding assay to determine apparent (fraction) or real (compound) IC<sub>50</sub>*

### Analysis of Overt Toxicity Due to Veraguamide C in Mice

Our cytotoxicity data above shows minimal effects of the veraguamides on non-TNBCs. These data suggest that the veraguamides would generally be safe and not exert overt toxicity in naïve animals. To test this hypothesis, a limited pilot blinded experiment was completed looking at locomotor behavior of mice following an intracerebroventricular (ICV) injection of 20 µg (0.5 µL volume) of veraguamide C compared to vehicle. In a 10-minute open field trial, there was no effect of veraguamide C on total distance traveled compared to vehicle (veraguamide C 35 m ± 4 sd n=4 vs control 31 m ± 10 sd n=4; unpaired t-test P>0.05).

These results are a promising starting point for the development of veraguamide analogs as non-toxic sigma receptor targeting agents. There has been considerable interest in the development of sigma 2/TMEM97 targeting agents for pain,^24^ neurodegeneration,^25^ antidepressant,^26^ and substance use disorder.^27^ Ongoing efforts to design, synthesize, and test analogs of these unique natural products are underway.

## Conclusions

In this manuscript we described the isolation of two new linear veraguamides, veraguamide O (1) and P (2), as well as the re-isolation of a known veraguamide, veraguamide C (3) from a Panamanian marine cyanobacteria. We tested the hypothesis that the linear depsipeptide veraguamide O (1) and cyclic veraguamide C (3) would be cytotoxic against cancer cell lines including TNBC lines that had not been assessed for any veraguamide compound. Surprisingly, while we confirmed mild toxicity of veraguamide C in ER positive cancer cells, we failed to find any evidence for toxicity of veraguamide O in these cells. This was in contrast to TNBC cells where both veraguamide C and O induced similar, albeit minor, levels of toxicity. This is the first example of veraguamides tested in TNBC cells. Overall these data suggest that the veraguamides are better suited for development as a non-toxic therapeutic and not for TNBC or other cancers. Molecularly, the veraguamides here show affinity for the sigma 2/TMEM97 receptor. We saw no overall toxicity seen in HEK-293 normal cells or in mice locomotor behavior. Thus, these compounds may also be valuable medicinal chemistry starting material for modulation of sigma 2/TMEM97 outside of cancer including in neurodegeneration^25^ and pain^7, 24, 26^.

## Materials and Methods

### General Experimental Procedures

NMR spectra were recorded with CD_3_CN (δ_C_ 118.2, δ_H_ 1.96), MeOD (δ_C_ 49.0, δ_H_ 3.31) and CDCl_3_ (δ_C_ 77.2, δ_H_ 7.26) as internal standards, on a Bruker 500 MHz spectrometer operating at 499.7 MHz for ^1^H and 125.7 MHz for ^13^C equipped with a 5 mm PATXI ^1^HD/D-^13^C/^15^N Z-GRD Probe. High-resolution mass measurements were obtained on an Agilent Technologies QTOF Accurate Mass mass detector. Additional mass spectra were acquired with benchtop Advion mass spectrometer. HPLC separation was carried out using a Dionex Ultimate 3000 pump system with UV detection using HPLC grade solvent. Accelerated chromatographic isolation was carried out on a Biotage Isolera One system with UV detection using HPLC grade solvent. Column chromatography was performed using Sorbent Technologies silica gel (230-400 mesh). Solvents were evaporated on a Heidolph rotary evaporator.

### Collection and Microscopy

Green/grey filamentous cyanobacteria were collected by hand on January 29, 2013 while snorkeling at a depth of ∼ 1 m off the coast of Isla Mina (GPS coordinates: N 8 29.717 W 78 59.947) in the Las Perlas Archipelago, Panama. The sample was fully suspended in 800 mL of 50:50 ethanol:seawater mixture to preserve the bacteria until it could be stored in a light-tight lab at -20 °C prior to extraction. A smaller sample was saved in RNAlater (ThermoFisher) for phylogenetic analysis. A voucher specimen (PLP-29Jan13-3) is deposited in the Department of Medicinal Chemistry, Graduate School of Pharmacy, Duquesne University.

Samples of DUQ0008 used for microscopy were prepared using a < 1 mm^3^ piece from the RNAlater sample. Filaments were spread out onto a slide with distilled H_2_O and imaged at 10X on a (Nikon 1AR-HD confocal microscope) using the FITC (487 nm), TRITC (560nm) and Cy5 (637 nm) and/or bright-field channels.

### 16S rRNA Gene Sequencing and Phylogenetic Analysis

Genomic DNA was extracted from DUQ0008 that was preserved in RNAlater after collection. Lysis buffer (10 mM Tris, 0.1 M EDTA, 0.5% (w/v) SDS, 20 μg/mL RNase, pH 8.0) was added to the cyanobacteria mass at 10X the biomass of the cells (i.e. 100 mg cells = 1000 μL lysis buffer). This was followed by 100 μL of lysozyme solution (10 mg/mL, Sigma) for 30 minutes with an incubation at 37°C. 0.01X the volume of Proteinase K (10 mg/mL, Gene Link) was then added and incubated for 1 hr at 50°C. Following this, the mixture was centrifuged at 13,000 rpm for 3 min and the remaining pellet was used with the Wizard Genomic DNA Purification Kit (Promega). Extracted genomic DNA then underwent PCR to amplify the 16S rRNA gene with primers (CYA106F and CYA1509R) previously utilized.^28, 29^ Successful PCR amplifications (∼1370 bp) underwent PCR purification using the Min Elute PCR Purification Kit (Qiagen) and purified PCR products were used for TOPO cloning. One Shot *E. coli* cells along with the pCR® 4-TOPO vector with kanamycin resistance from the TOPO TA Cloning ® Kit (Invitrogen) were used and cells were plated overnight at 37°C on LB KAN plates. Successful transformants were selected and used in colony PCR with the M13F and M13R primers to confirm the presence of the 16S rRNA sequence now in the TOPO vector (∼1535 bp). Purified PCR product for correct samples was sent to Beckman Coulter (now GENEWIZ) to undergo forward and reverse Sanger sequencing with the M13F and M13R primers. The sequence is available in the GenBank database under the accession number MH345835. The sequence for DUQ0008, along with 16S rRNA gene sequences that were obtained with sequence data available at the National Center for Biotechnology (NCBI) webpage (http://www.ncbi.nlm.nih.gov) were then used for phylogenetic analysis. DUQ0008 gene sequence was first aligned in an unbiased fashion using the NCBI Standard Nucleotide BLAST^18^ to determine the closest alignment. Five of the top six alignment hits were from the newly characterized *Okeania* genus.^19^ We next aligned DUQ0008 with the top NCBI BLAST *Okeania* hit and all *Okeania* type gene sequences from a recent phylogenetic analysis of natural product-producing cyanobacteria.^21^ Sequences were aligned using the Muscle algorithm and phylogeny reconstruction was done using maximum likelihood method with 600 bootstrap replicates using the MEGA (version 5.1) program.^30^

### Extraction and Isolation

The cyanobacterial biomass (75 g, dry wt) was extracted exhaustively with 2:1 CH_2_Cl_2_–MeOH to afford 3.4 g of crude extract. This crude extract was fractionated over normal phase silica gel with a stepwise gradient solvent system of increasing polarity starting from 100% hexanes to 100% MeOH, to yield nine fractions (A–I). The fraction eluting with 100% MeOH (fraction I, 150 mg) was chromatographed over Hypersep C18 (1000 mg, 8 mL) cartridge, and the column eluted with 30 mL of 50% and 75% MeOH–water mixture, then 100% MeOH to yield sub-fractions I1–3 corresponding to 50.7, 73.1 and 13.9 mg respectively. A portion of the fraction eluted with 50% acetonitrile in water (sub-fraction I1, 30.3 mg) was subjected to accelerated reversed-phase (Biotage, Snap KP-C18-HS, 12 g cartridge) chromatography using a linear gradient of acetonitrile–water (50% acetonitrile in 9 CV followed by 50–100% acetonitrile in 8 CV, and then 100% acetonitrile for 9 CV). Combinations of 22 fractions, A–E, were obtained based upon peaks observed in the chromatogram generated from UV detection at 210 and 254 nM. Sub-fraction D (9.4 mg) was subjected to reversed-phase HPLC (Phenomenex, Synergi-fusion 4μ, 150 x 10 mm) using a linear gradient of acetonitrile–H_2_O acetonitrile–0.1%TFA (in water) gradient (55% acetonitrile for 22 min, then 55–100 % acetonitrile in 5 min and then 100% acetonitrile for 5 min) to yield veraguamide O (**1**) (2.3 mg, 14.24 mins) and veraguamide P (**2**) (0.3 mg, 11.17 mins). Sub-fraction E which was comprised of a mixture of **1** and **2** (3.9 mg), as confirmed by ^1^H NMR and mass spectral analysis, was chromatographed under similar conditions to afford **1** (1.5 mg, 14.33 mins) and **2** (0.5 mg, 11.25 mins). Fraction G (564.8 mg) was subjected to silica gel column chromatography and yielded seven fractions, G1–7. Sub-fraction G1 (52.3 mg) was further chromatographed over reversed-phase HPLC (Phenomenex, Synergi-fusion 4μ, 150 x 10 mm; 83% MeOH:H_2_O for 20 min, followed by 100% MeOH for 10 min; 1.5 mL/min) to afford a sub-fraction (12.0 mg), collected between 25.0–30.0 min. This sub-fraction was subjected to accelerated reversed-phase (Biotage, Snap KP-C18-HS, 12 g cartridge) chromatography using a linear gradient of methanol-water (50% MeOH in 2.5 CV then 50-100% in 16 CV; 30 mL/min). The fraction collected at 10 CV afforded veraguamide C (**3**, 1.7 mg) after concentration in vacuo.

*Veraguamide O* (***1***): colourless oil; ^1^H-,^13^C- and 2D-NMR see Table 1; HRESIMS m/z 773.4361 [M + K] (calcd. for C_39_H_66_N_4_O_9_K, 773.4466), m/z 757.4626 [M + Na] (calcd. for C_39_H_66_N_4_O_9_Na, 757.4727) and m/z 735.4804 [M + H] (calcd. for C_39_H_67_N_4_O_9,_ 735.4908_)_.

*Veraguamide P* (***2***): colourless oil; ^1^H-,^13^C- and 2D-NMR see Table 1; HRESIMS m/z 743.4626 [M + Na] (calcd. for C_38_H_64_N_4_O_9_Na, 743.4571).

*Veraguamide C* (***3***): colourless oil; ^1^H-NMR – comparable to previously reported literature values. ESIMSm/z 733.98 calcd. for C_37_H_95_N_4_O_9_Na_2_ [M + 2Na -H].

### Methanolysis/Partial Hydrolysis of 1 and 2

A sample of **1** (0.5 mg) was treated with 5% NaOH (v/v) in methanol at room temperature for 24 h. The reaction mixture was concentrated to dryness, and the residue was partitioned between H_2_O and EtOAc. The organic layer was collected and dried and the identity of the crude methanolysis products was confirmed by ESI-MS. A similar procedure was repeated for **2**.

### Cell Culture

The human triple-negative breast cancer cell line MDA-MB-231 was obtained from ATCC and was maintained in DMEM:F-12 (1:1) (Life Technologies), 10% Fetal Bovine Serum (Atlanta Biologicals), and 1% Penicillin/Streptomycin (Sigma). The estrogen positive MCF-7 cell line was also obtained from ATCC and was maintained in 1640 RPMI media (Life technologies), 10% FBS, and 1% Penicillin/Streptomycin. HEK-293 cells were also obtained from ATCC and maintained in Dulbecco’s Modification of Eagle’s Medium (DMEM), 10% Fetal Bovine Serum (FBS) and 1% Antibiotics Penicillin and Streptomycin (Pen/Strep).

### Cytotoxicity Assays

Two rounds of toxicity experimentation were completed. First, MDA-MB-231 and MCF-7 cells were plated at a density of 5×10^3^ cells per well in a 96 well plate. Cells were allowed to attach overnight. The cells were treated with depsipeptides or vehicle for 72 hours with 5% FBS stimulation. DMSO was used as a vehicle control. After treatment, 10 µL MTT (Sigma) was added to each well (0.5 mg/mL final concentration) and the plates were incubated for 3 hours (5% CO_2_ and 37(). The medium was removed and the MTT-formazan crystals were dissolved with 100 µL DMSO per well. The absorbance was measured at 570 nm with a VICTOR^3^ 1420 multilabel counter (Perkin Elmer). Three wells were analyzed for each condition, and wells containing medium-MTT only (no cells) and medium-MTT (DMSO + 5% FBS) were used as controls. Results were normalized to the DMSO + 5% FBS group.

Second, HEK-293 cells were plated at a density of 3×10^4^ cells per well in a 96-well plate. Cells were allowed to attach overnight. The cells were treated with veraguamides C and P (or vehicle 0.5% DMSO). Veraguamides were suspended in DMEM to achieve final concentrations ranging from 0.01 µM to 10 µM in 0.5% DMSO (final concentration in well). 11 replicate wells were used for each concentration of each treatment. Cells were treated for 24 hours. After 24 hours of exposure to the test compounds, cell viability was assessed using the CellTiter-Blue® assay. Briefly, 20 µL of CellTiter-Blue® reagent was added directly to each well containing 100 µL of media. The plates were then incubated for an additional 4 hours at 37 °C to allow for sufficient metabolic conversion. The fluorescence was measured using a fluorescence microplate reader set to an excitation wavelength of 560 nm and an emission wavelength of 590 nm. The percentage of viable cells was calculated by comparing the fluorescence readings of the treated wells (11 per concentration) with those of the control DMSO wells.

### Psychoactive Drug Screening Program (PDSP) Receptor Screening

All binding data for fractions and compounds were performed and generously provided by the National Institute of Mental Health’s Psychoactive Drug Screening Program (NIMH PDSP) utilizing a radioligand competition-binding assay. Experimental details are available online at http://pdsp.med.unc.edu/. In brief, fractions that pass primary binding criterion of greater than 50% inhibition at 10 μM are tagged for secondary radioligand binding assay to determine the apparent or real IC_50_. Data reported here are for affinity of fractions and compounds to the sigma receptors.

### In Vivo Testing

All animal procedures were reviewed and carried out in accordance with the National Institutes of Health Guide for the Care and Use of Animals and the Institutional Animal Care and Use Committee at Duquesne University. Experiments were performed on male C57Bl/6J mice that were 8–10 weeks old.

#### Surgical Procedures

Animals were anesthetized with 3% isoflurane/0.6% O_2_ during all surgical procedures. Intracerebroventricular (ICV) cannulation surgeries were performed as described previously.^31^ Briefly, mice were placed in a stereotaxic frame and an 8.00 mm steel cannula was placed into the right lateral ventricle at the following anatomical coordinates: 0.5 mm anterior to bregma, 1.0 mm lateral to midline and 2.0 mm ventral to the skull. A dental cement skullcap secured with two bone screws was used to hold the cannula in place. Mice recovered on heating pads and were given one week of recovery prior to the beginning of behavioral testing. Following behavioral testing, cannula placement was verified with necropsy.

#### Open Field

Open field was completed as previously described.^32^ Briefly, mice were habituated to a dimly lit room for 1 hour with 60dB white noise. Veraguamide C was dissolved in a mixture of 50% DMSO and 50% artificial cerebrospinal fluid (aCSF) containing the following (in mM): 125 NaCl, 2.5 KCl, 1.25 NaH_2_PO_4_, 25 NaHCO_3_, 2.0 CaCl_2_, 1.0 MgCl_2_ and 25 d-glucose, bubbled with 95% O_2_/5% CO_2_ for 20 minutes prior to use. Microinjections were performed using a 32-gauge injector that extended 0.5 mm beyond the tip of the ICV cannula. The injector was attached to flexible tubing and a 1.0 μL syringe (Hamilton) that was used to deliver a total volume of 0.5 μL over a 2-minute period. The injector was kept in place for an additional 1 minute to allow complete compound infusion. Veraguamide C was administered at a 20 μg dose. 50%/50% aCSF/DMSO (0.5 μL) was used as a vehicle control. Mice were treated with veraguamide C or vehicle 5 minutes before testing in the open field box (40.6 x 40.6 x 30cm). Behavior was videorecorded and scored using ANY-Maze software (Stoelting Co. Version 4.98). Total distance travelled (m) was measured. Injection and analysis was completed blinded to treatment.

## ACKNOWLEDGEMENTS

We would like to thank Analise Zapadka, Edward Hilton, and Youstina Seliman for technical assistance. We thank the following departments, funding agency and sponsors: Autoridad Nacional del Ambiente de Panamá (ANAM) and the Smithsonian Tropical Research Institute (STRI); NIH NCCIH R15AT008060 (BJK, KJT); NIH NINDS R61NS127271 (KJT, BJK); Fogarty International Center Panama International Cooperative Biodiversity Grant TW006634 (partially supported collection of material by KJT). This work was further supported by the Center for Pharmaceutical Research and Innovation (CPRI, NIH P20 GM130456).

## SUPPORTING INFORMATION

Supporting information contains ^1^H NMR, COSY, TOCSY, HSQC, HMBC, and HRESIMS spectra for **1** and **2**.

## Notes

### Competing Interest Statement

The authors have declared no competing interest.

