## Supplemental Figures for "Isolation and Bioassay of Linear Veraguamides from a Marine Cyanobacterium (*Okeania sp.*)"

Table of Contents:

S 1: 1H-NMR spectrum of natural product **1** in CD_3_CN at 500 MHz.	pg 5

S 2: COSY-NMR spectrum of natural product **1** in CD_3_CN at 500 MHz.	pg 6

S 3: TOCSY-NMR spectrum of natural product **1** in CD_3_CN at 500 MHz.	pg 7

S 4: HSQC-NMR spectrum of natural product **1** in CD_3_CN at 500 MHz.	pg 8

S 5: HMBC-NMR spectrum of natural product **1** in CD_3_CN at 500 MHz.	pg 9

S 6: HRES-Mass spectrum of natural product **1** (positive ionization mode).	pg 10

S 7: 1H-NMR spectrum of natural product **2** in CD_3_CN at 500 MHz.	pg 11

S 8: COSY-NMR spectrum of natural product **2** in CD_3_CN at 500 MHz.	pg 12

S 9: HSQC-NMR spectrum of natural product **2** in CD_3_CN at 500 MHz.	pg 13

S 10: HMBC-NMR spectrum of natural product **2** in CD_3_CN at 500 MHz.	pg 14

S 11: HRES-Mass spectrum of natural product **2** (positive ionization mode).	pg 15

S12: 1H-NMR spectrum of natural 2 in CD3OD at 500 MHz	pg 16

S13: 1H-NMR spectrum of natural 3 in XXX at 500 MHz	pg 17

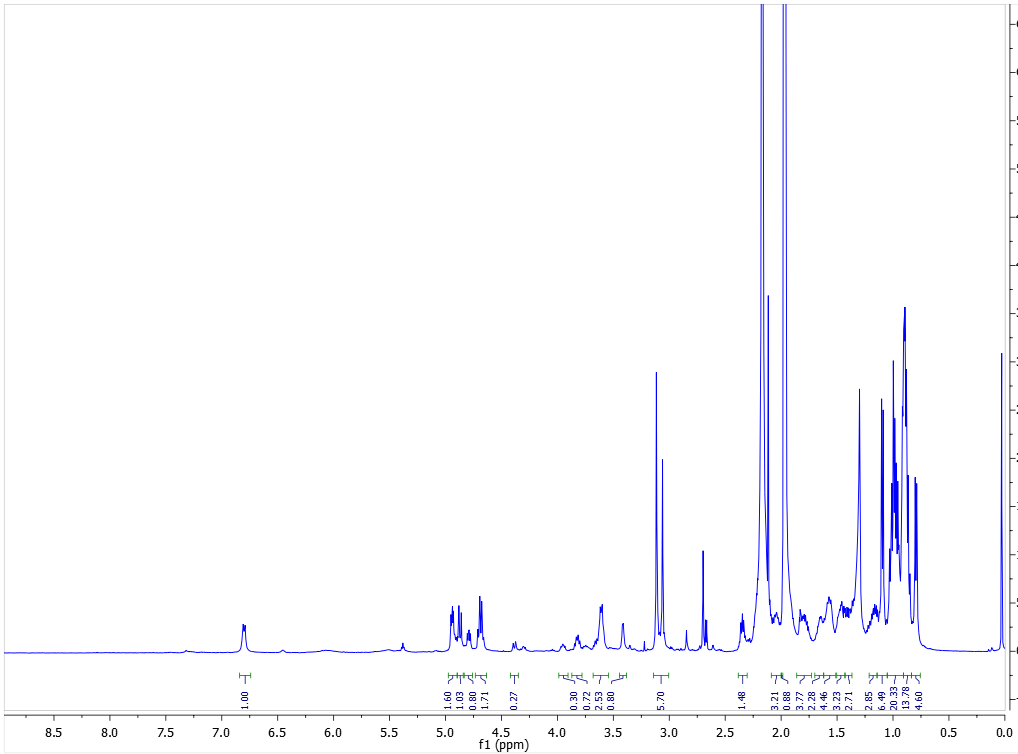
Figure **S1**. ^1^H NMR spectrum of compound (**1**) in CD_3_CN at 500 MHz.

**Figure S2**. COSY spectrum of compound (**1**) in CD_3_CN at 500 MHz.

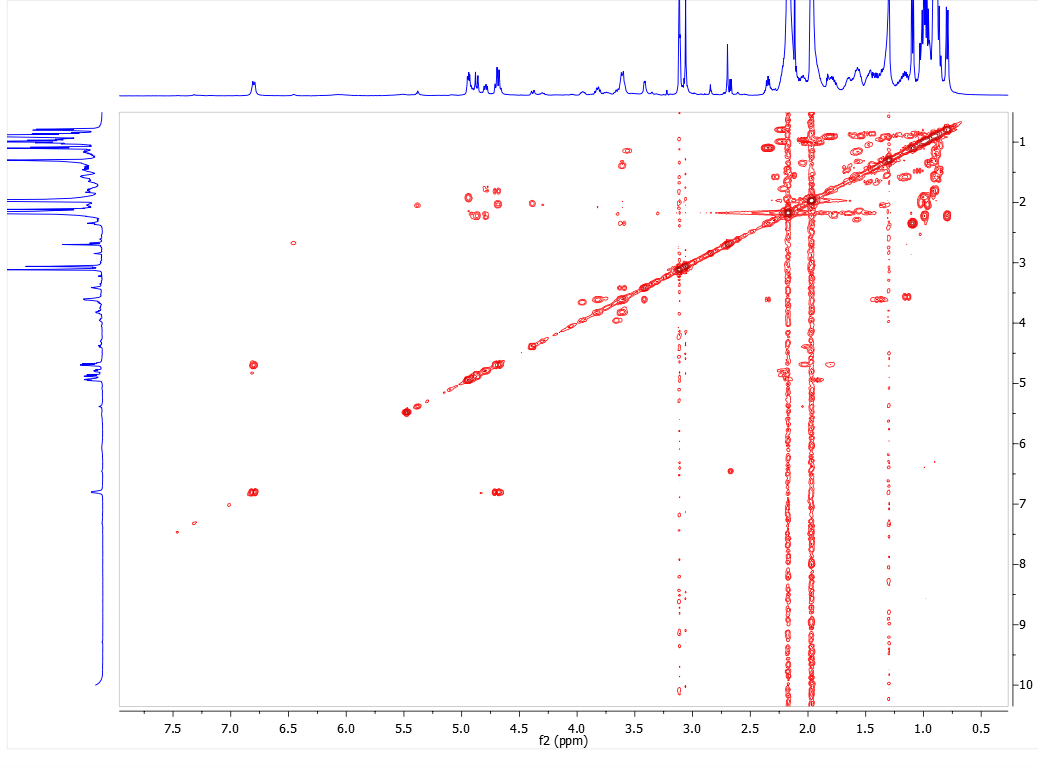

**Figure S3**. TOCSY spectrum of compound (**1**) in CD_3_CN at 500 MHz.

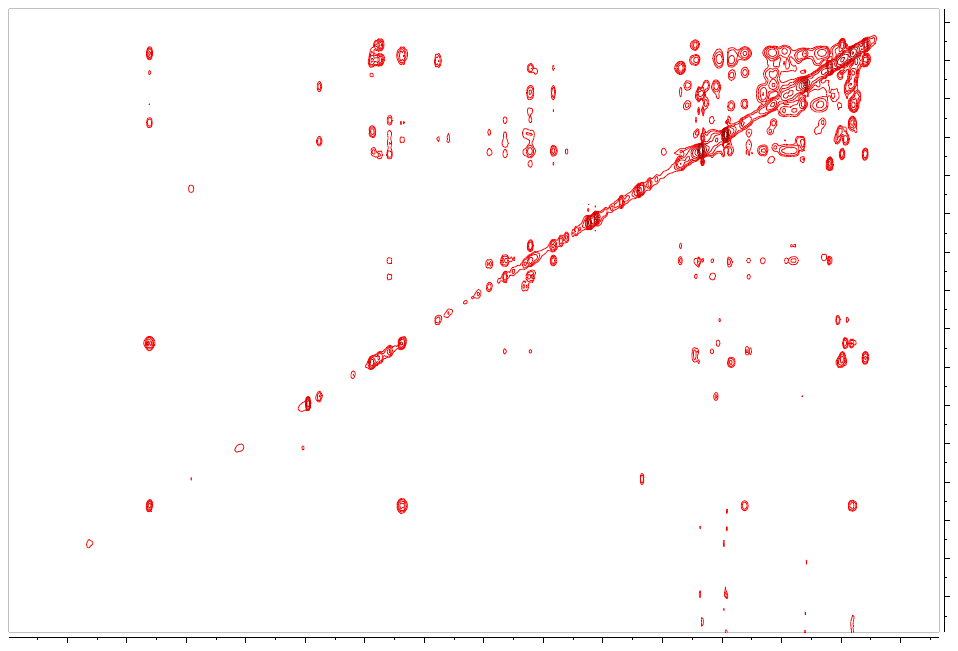

**Figure S4**. HSQC spectrum of compound (**1**) in CD_3_CN at 500 MHz.

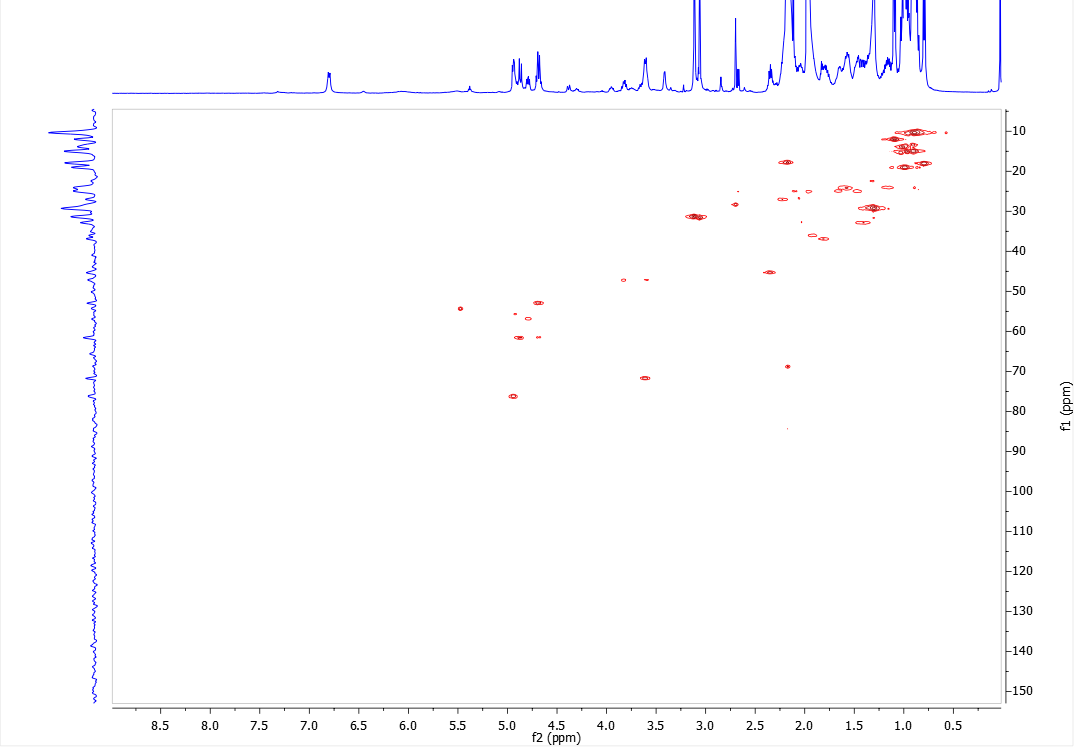

**Figure S5**. HMBC spectrum of compound (**1**) in CDCl_3_ at 500 MHz.

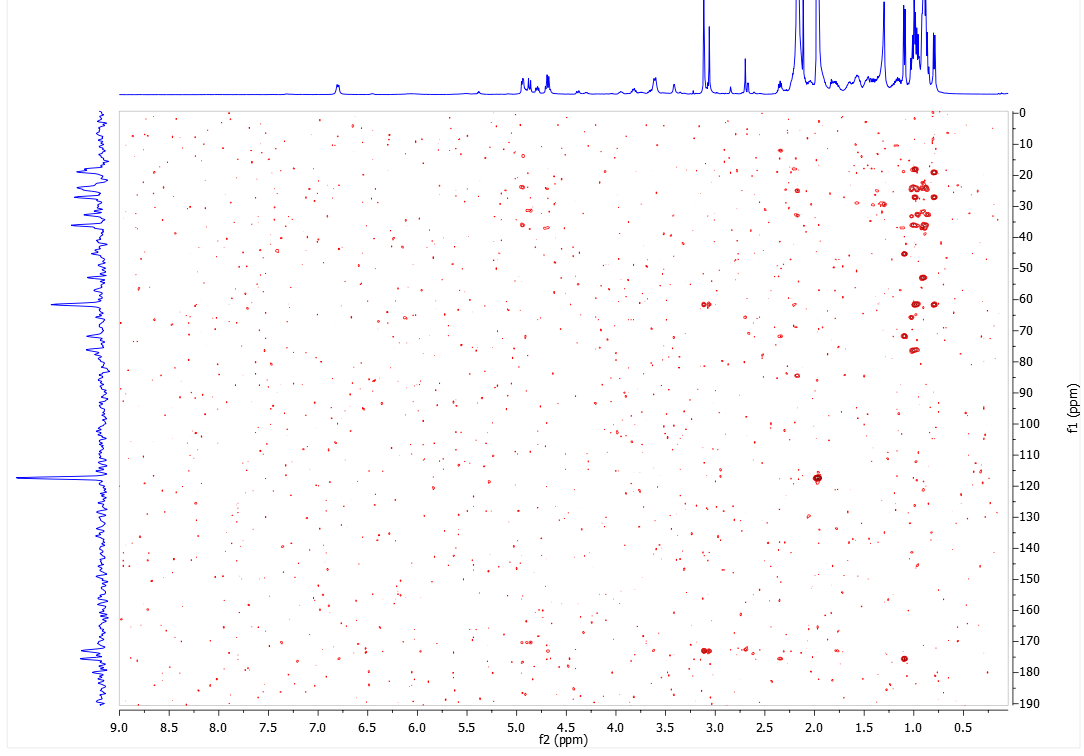

**Figure S6**. HRESIMS spectrum of (**1**).

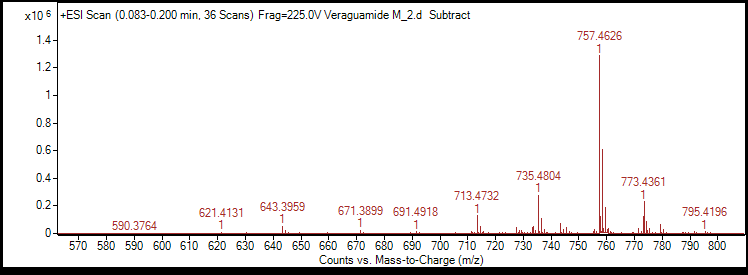

**S 7**: ^1^H-NMR spectrum of *natural product* **2** in CD_3_CN at 500 MH
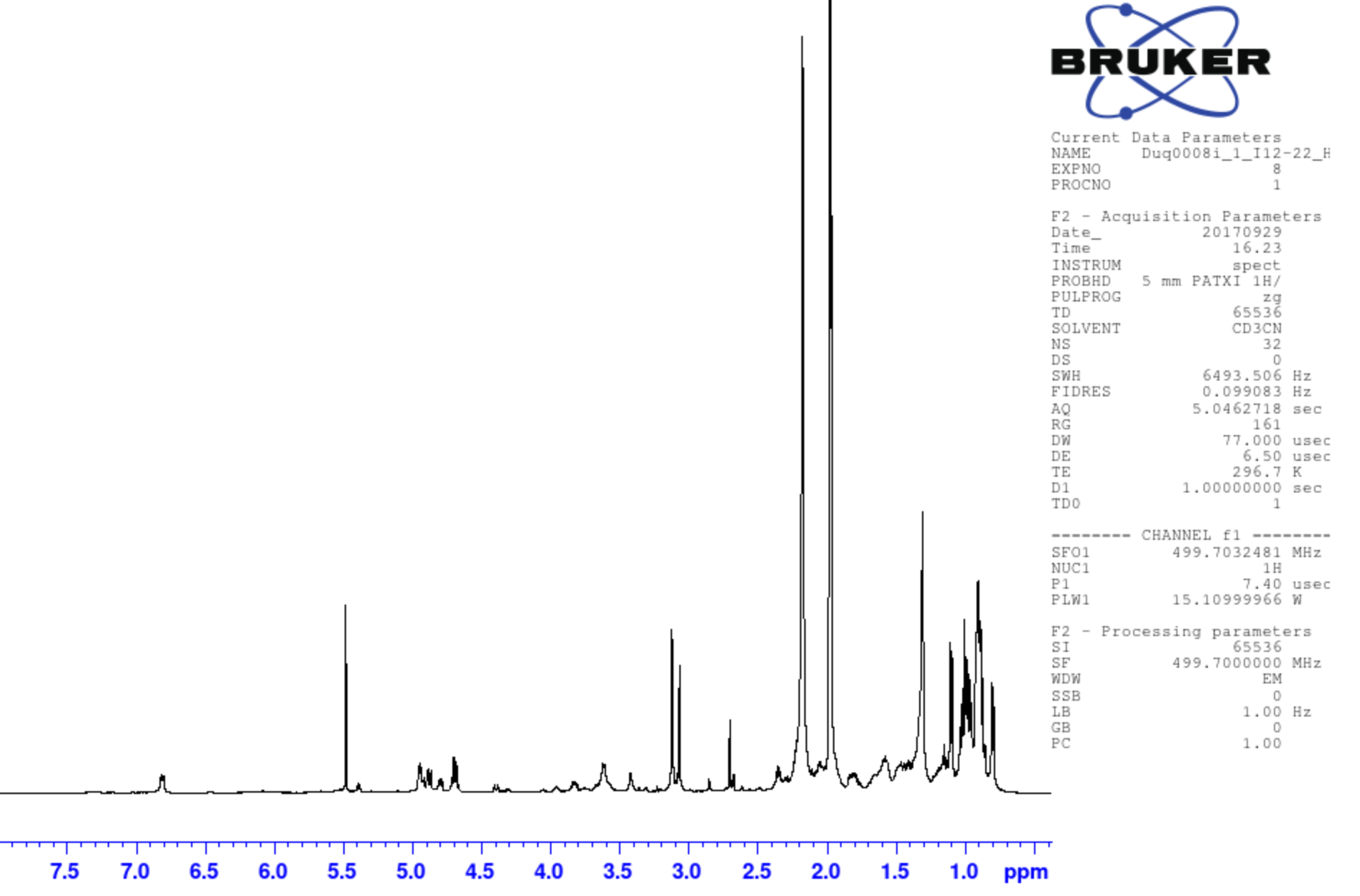

**S 8**: COSY-NMR spectrum of *natural product* **2** in CD_3_CN at 500 MHz.
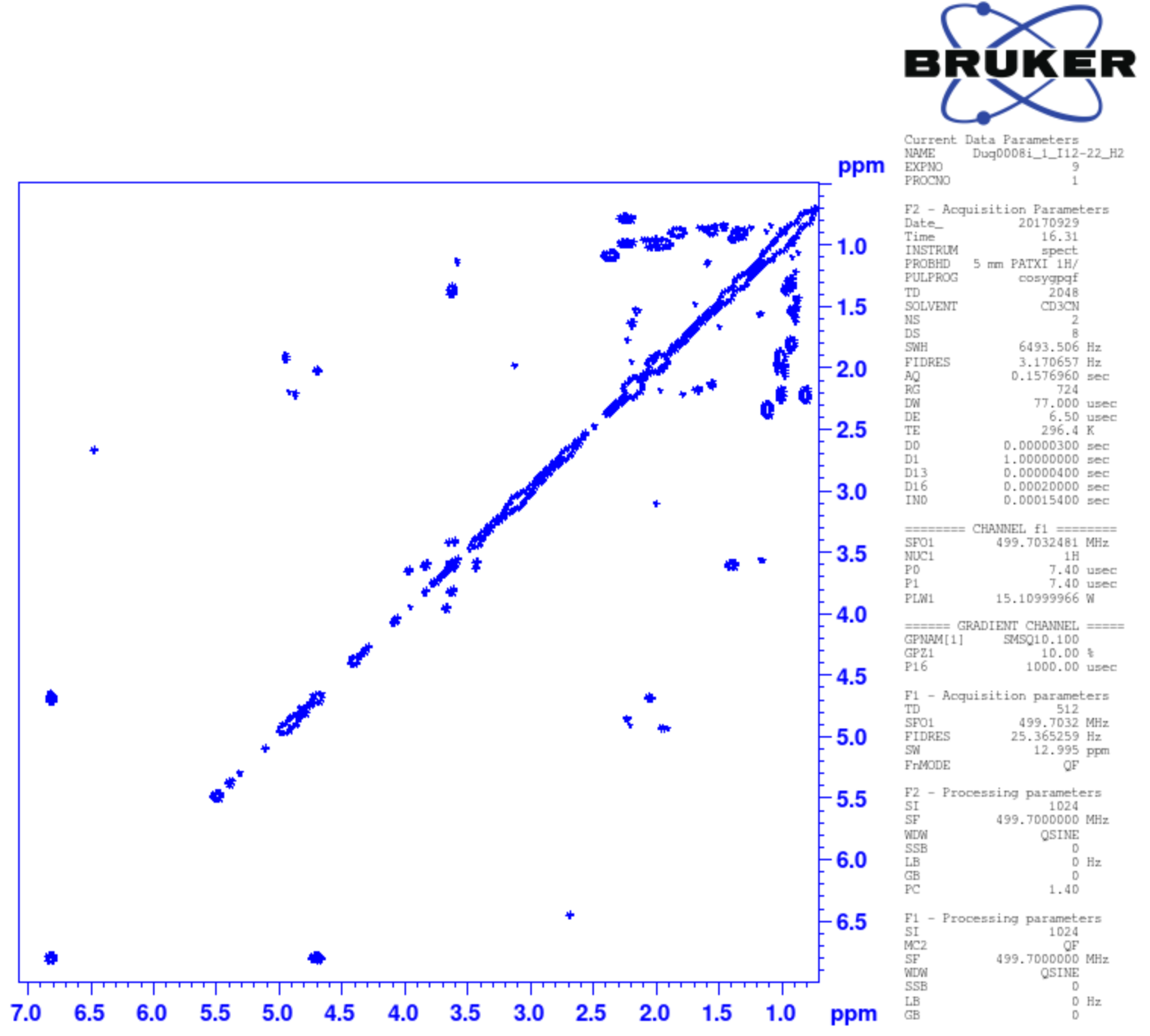

**S 9**: HSQC-NMR spectrum of *natural product* **2** in CD_3_CN at 500 MHz.

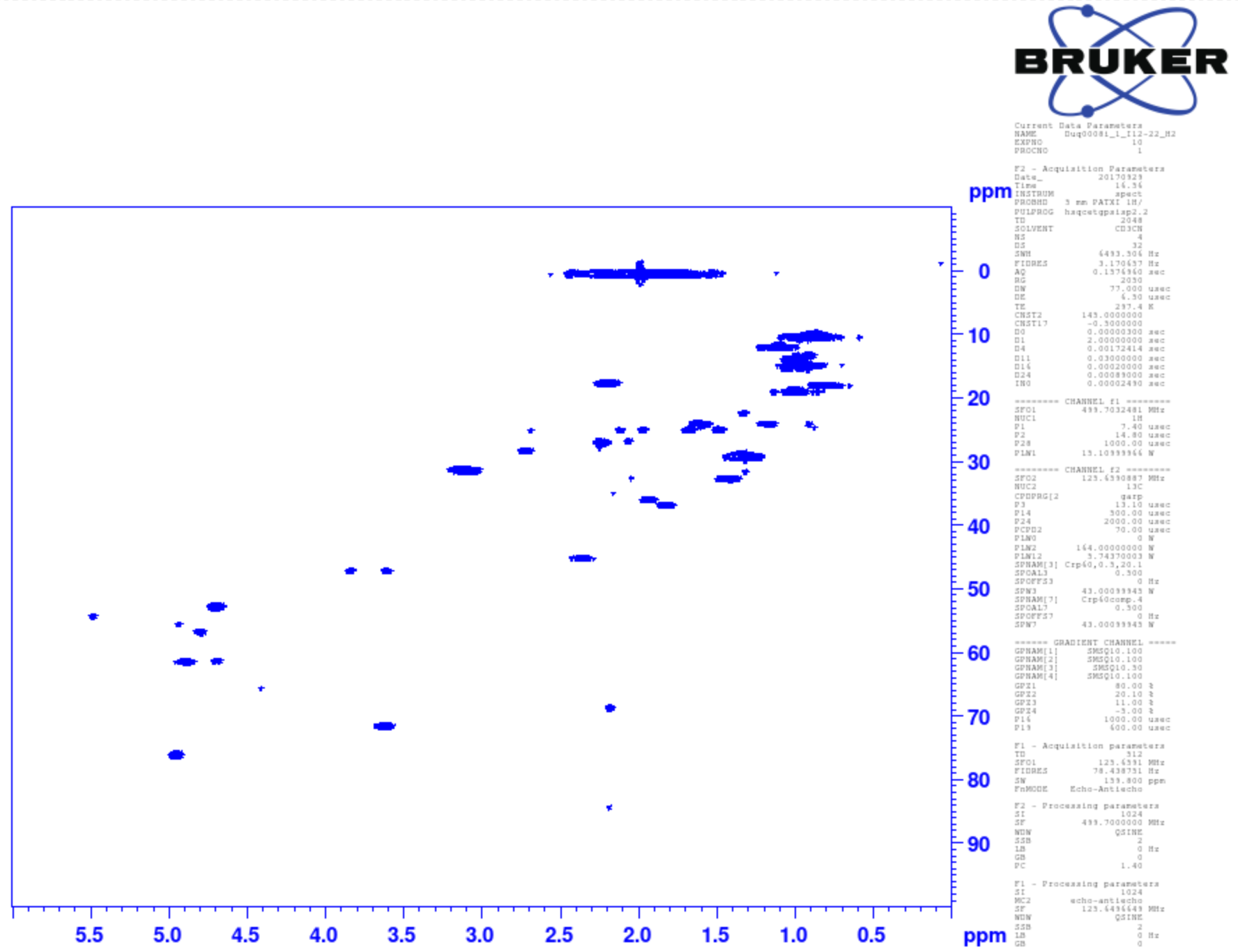

**S 10**: HMBC-NMR spectrum of *natural product* **2** in CD_3_CN at 500 MHz.

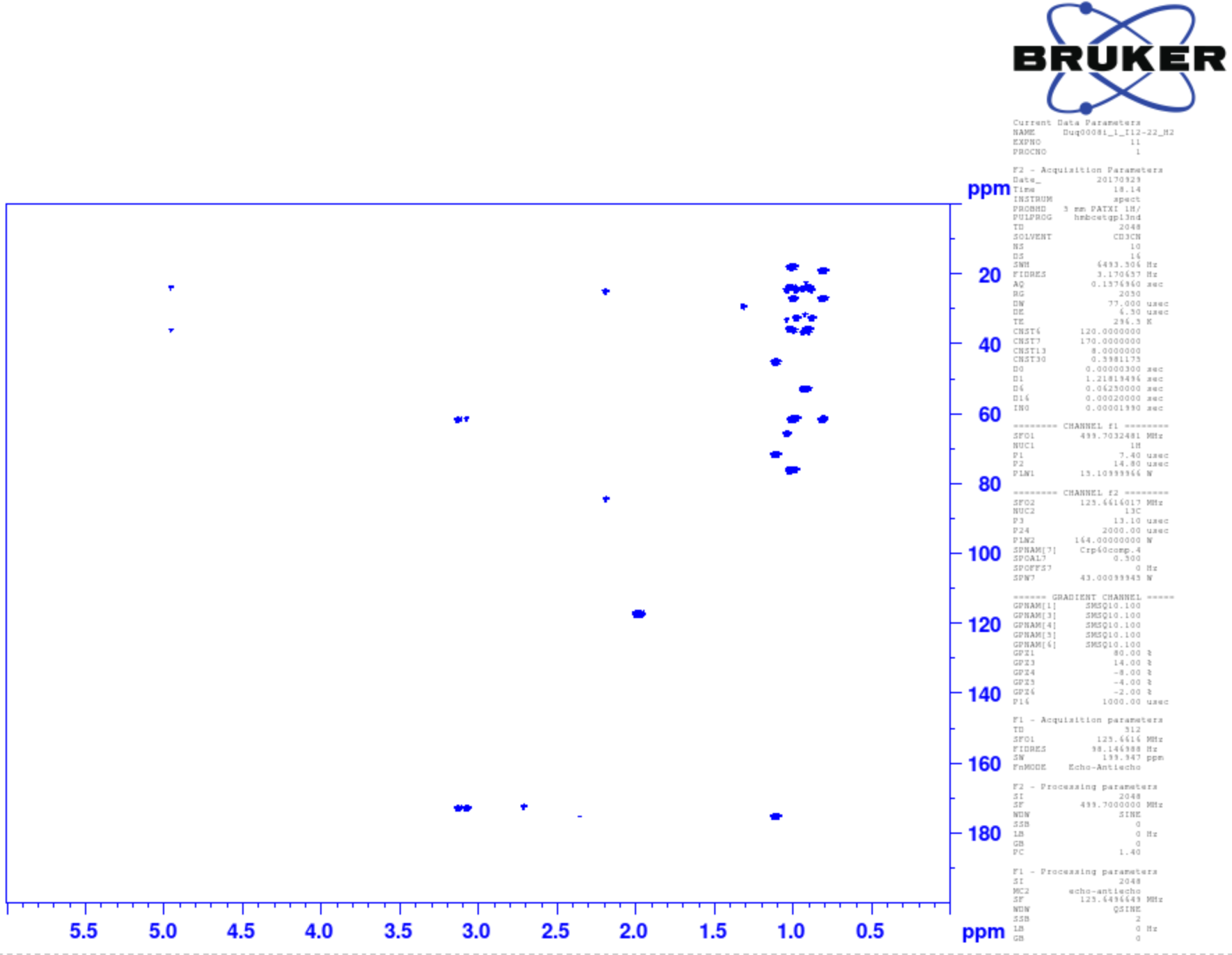

**S 11**: HRES-Mass spectrum of *natural product* **2** (positive ionization mode).

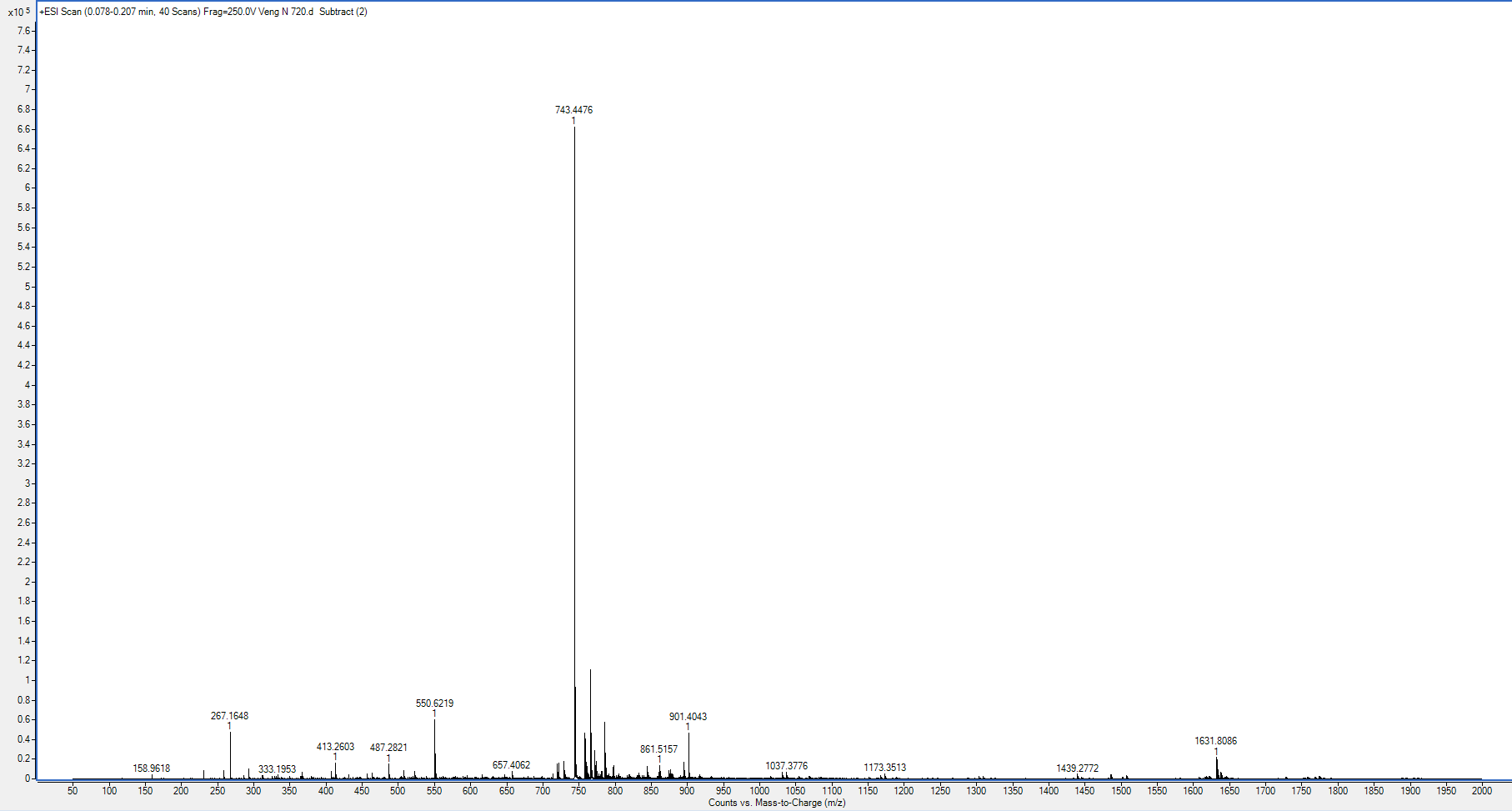

**S 12**: ^1^H-NMR spectrum of *natural product* **2** in CD_3_OD at 500 MHz.

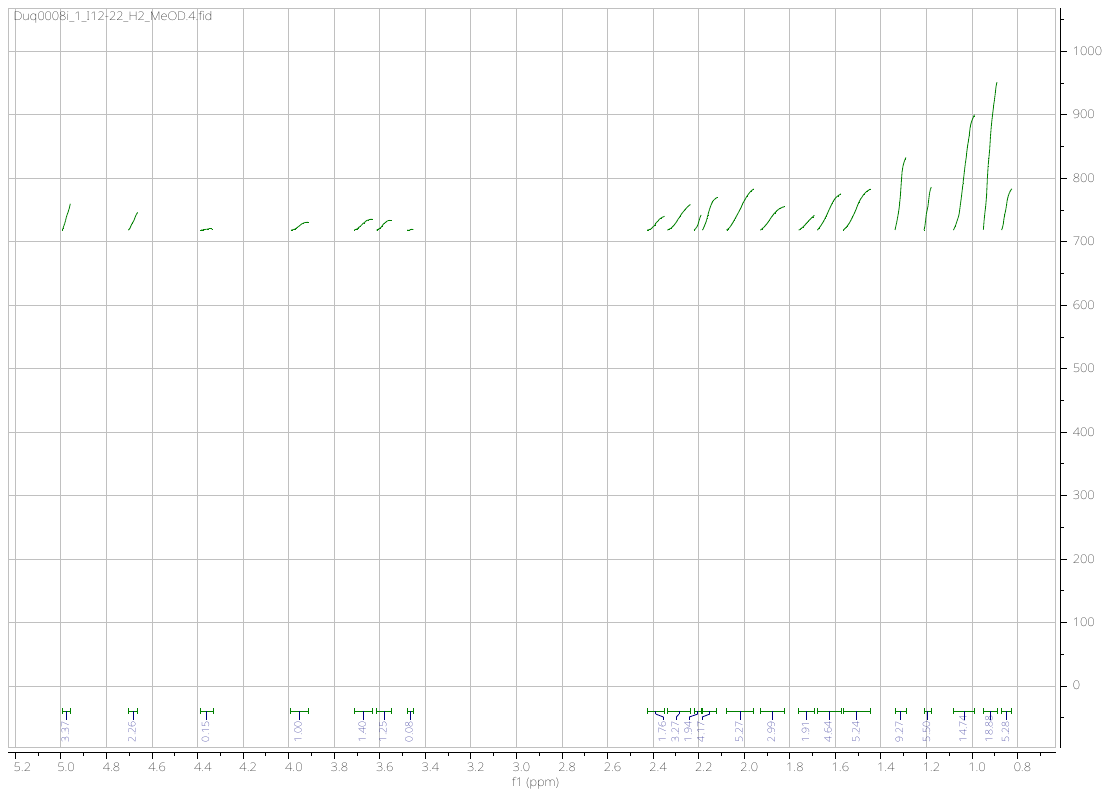
